# ^56^Fe ion exposure increases the incidence of lung and brain tumors at a similar rate in male and female mice

**DOI:** 10.1101/2023.06.06.543754

**Authors:** Sophie R. Finkelstein, Rutulkumar Patel, Katherine Deland, Joshua Mercer, Bryce Starr, Daniel Zhu, Hooney Min, Michael Reinsvold, Lorraine Da Silva Campos, Nerissa Williams, Lixia Luo, Yan Ma, Jadee Neff, Mark Hoenerhoff, Everett J. Moding, David G. Kirsch

## Abstract

The main deterrent to long-term space travel is the risk of Radiation Exposure Induced Death (REID). The National Aeronautics and Space Administration (NASA) has adopted Permissible Exposure Levels (PELs) to limit the probability of REID to 3% for the risk of death due to radiation-induced carcinogenesis. The most significant contributor to current REID estimates for astronauts is the risk of lung cancer. Recently updated lung cancer estimates from Japan’s atomic bomb survivors showed that the excess relative risk of lung cancer by age 70 is roughly four-fold higher in females compared to males. However, whether sex differences may impact the risk of lung cancer due to exposure to high charge and energy (HZE) radiation is not well studied. Thus, to evaluate the impact of sex differences on the risk of solid cancer development post-HZE radiation exposure, we irradiated *Rb^fl/fl^; Trp53^fl/+^* male and female mice infected with Adeno-Cre with various doses of 320 kVp X-rays or 600 MeV/n ^56^Fe ions and monitored them for any radiation-induced malignancies. We observed that lung adenomas/carcinomas and esthesioneuroblastomas (ENBs) were the most common primary malignancies in X-ray and ^56^Fe ion-exposed mice, respectively. In addition, 1 Gy ^56^Fe ion exposure compared to X-rays led to a significantly higher incidence of lung adenomas/carcinomas (p=0.02) and ENBs (p<0.0001). However, we did not find a significantly higher incidence of any solid malignancies in female mice as compared to male mice, regardless of radiation quality. Furthermore, gene expression analysis of ENBs suggested a distinct gene expression pattern with similar hallmark pathways altered, such as MYC targets and MTORC1 signaling, in X-ray and ^56^Fe ion-induced ENBs. Thus, our data revealed that ^56^Fe ion exposure significantly accelerated the development of lung adenomas/carcinomas and ENBs compared to X-rays, but the rate of solid malignancies was similar between male and female mice, regardless of radiation quality.

## Introduction

The National Aeronautics and Space Administration (NASA) plans to send astronauts back to the Moon and, eventually, on the first trip to Mars by the end of this decade. However, many risk factors are associated with long-term space travel, including exposure to space radiation from galactic cosmic rays (GCRs) and solar particle events (SPEs) contributing to the risk of Radiation Exposure Induced Death (REID) [1, 2]. Although protons and helium ions comprise the majority of space radiation, high charge and energy (HZE) radiation from GCRs and SPEs have high linear energy transfer (high-LET), leading to clustered damage in tissues that may be particularly carcinogenic [3]. NASA has kept the REID limit to 3% for the risk of death due to radiation-induced carcinogenesis [4], but this risk assessment has large uncertainties and is estimated based on data for radiation carcinogenesis from radiation exposure on Earth. In fact, the risk of radiation-induced lung cancer is the largest contributor to current REID estimates for astronauts. One of the best data sets on radiation exposure and human cancer risk comes from atomic bomb survivors. The Radiation Effects Research Foundation in Hiroshima, Japan, estimates that following radiation exposure at the age of 30, the risk of solid cancer by the age of 70 increases by 35% per Gy for men and 58% per Gy for women [5]. Recent updates for lung cancer show that for a 30-year-old, never-smoker male and female exposed to radiation, the excess relative risk of lung cancer by age 70 is 0.34 per Gy and 1.32 per Gy (i.e., 3.91 fold higher), respectively [6]. The clear sex dependence of lung cancer in humans after radiation exposure forms the basis for risk assessment models where female astronauts have nearly twice the estimated REID as male astronauts. However, whether HZE ions pose an even higher risk of lung cancer in female astronauts remains to be studied using animal models.

Prior studies of radiation carcinogenesis for lung cancer have utilized specific inbred strains of “wild-type” mice [7, 8] or genetically engineered mouse models (GEMMs) [9, 10]. Although some may argue that “wild-type” strains of mice are a more relevant model for human astronauts exposed to radiation, “wild-type” mice do not model multi-step carcinogenesis that results from accumulated mutations in a healthy individual over time. Most astronauts will be in the middle-aged group when they embark on a long-term mission to Mars. Thus, some but not all cells within the lung will already harbor mutations in key tumor suppressor genes due to DNA replication error [11-13]. The tumor suppressors, such as *TP53* and *RB*, are frequently inactivated via deletion or inactivation through promotor hypermethylation in most human lung cancers [14-16]. Here, we utilized a conditional mouse model in which alleles of *Trp53* and *Rb* can be deleted following the delivery of an adenovirus expressing Cre-recombinase to the lungs to determine whether X-ray ionizing radiation (low-LET IR) or HZE ion exposure (high-LET IR) and sex differences affect lung tumor development.

## Materials and Methods

### Animals

Institutional Animal Care and Use Committee-approved protocols were followed at Duke University and Brookhaven National Laboratory (BNL) for all animal experiments. *Rb^fl/fl^; Trp53^fl/+^; Rosa26^LSL-YFP/+^* (RPY) mice were generated using the stepwise breeding of *Rb1^tm2Brn^*/J (*Rb^fl/fl^*) [17], B6.129P2-*Trp53^tm1Brn^*/J (*Trp53^fl/+^*) [17], and B6.129X1-*Gt(ROSA)26Sor^tm1(EYFP)Cos^*/J (*Rosa26^LSL-YFP/+^*) [18] mice. The resulting mice were of mixed genetic backgrounds. In addition, these mice contained a single reporter allele in *Rosa26* locus in the form of lox-stop-lox (LSL)-YFP with no impact on tumorigenesis and, thus, they are referred to as *Rb^fl/fl^; Trp53^fl/+^*mice. To minimize the impact of genetic background, age-matched littermate controls were used so that the potential genetic modifiers would be randomized among all groups. All animals were bred at Duke University, where mice were exposed to whole-body irradiation with X-rays. Mice were shipped to BNL for whole-body exposure to ^56^Fe ions. All animals were kept in ventilated cages in a room with constant temperature (72 ± 2° F) and a 12:12h light:dark cycle at Duke University and BNL.

### Ad-Cre infection and Low-and high-LET radiation exposure

To delete both copies of *Rb* and 1 allele of *Trp53*, an adenovirus (Ad5CMV-Cre, University of Iowa Viral Vector Core) expressing Cre recombinase (Adeno-Cre) was introduced intranasally to *Rb^fl/fl^; Trp53^fl/+^*male and female mice at Duke University. Briefly, 25 μl of Adeno-Cre was added to 600 μl of DMEM media, followed by the addition of 3 μl of 2M CaCl_2_. Prepped Adeno-Cre solution was kept for 15 minutes. After a brief incubation, each mouse received 50 μl of prepped Adeno-Cre solution intranasally. Mice exposed to ^56^Fe ions were shipped to BNL. Exposure to radiation occurred 12-15 days after Adeno-Cre infection. On the day of high-LET radiation exposure, mice were put in 50 ml conical tubes, arranged in a foam holder, and placed in an upright position perpendicular to a 20×20 beam line. Unanesthetized mice were irradiated with 0.2, 0.5, or 1 Gy ^56^Fe 600 MeV/n ions (LET ∼ 175 KeV/ µm) at a dose rate of 0.2 Gy/min. For X-ray irradiations, male and female mice were exposed at Duke University using an X-RAD 320 Biological Irradiator (Precision X-Ray). Unanesthetized mice were arranged in a pie-shaped animal holder and irradiated with 1, 2, or 4 Gy at a dose rate of 2 Gy/min using 320 kVp X-rays. One cohort of sham-irradiated (0 Gy) mice were shipped to BNL, and another cohort of sham-irradiated (0 Gy) mice were kept at Duke. Both sham-irradiated cohorts were treated the same way as irradiated cohorts to minimize the impact of travel-related stress on experiment outcomes. As there was no difference in the time to tumor or overall survival between the two sham-irradiated cohorts, they were combined and compared with irradiated cohorts for the analysis.

### Necropsy and Histopathology analysis

Mice were monitored for tumor development and followed up to 2 years (1^st^ cohorts of mice) or up to 1 year (2^nd^ cohorts of mice) post-irradiation (IR). Any mice found moribund/sick during weekly checks were euthanized, and a complete necropsy was performed. Post complete necropsy, major organs, including the lungs, spleen, liver, thymus, cervical lymph nodes, brain, and any abnormal masses, were collected in 10% formalin. For each mouse, the lung was inflated with 10% formalin delivered by intratracheal injection for optimal preservation. All collected tissues fixed in 10% formalin overnight were transferred to 70% ethanol for short-term storage. Formalin-fixed tissues were processed, embedded in paraffin, sectioned, and stained using hematoxylin and eosin (H&E). A 5 µm-thick section was cut using each formalin-fixed paraffin-embedded (FFPE) tissue. For the lung, three sections per tissue were cut for H&E staining by removing 100 µm tissue between each section. Histopathology was performed by a board-certified veterinary pathologist, Dr. Mark Hoenerhoff, at the University of Michigan, Ann Arbor, MI.

### Mouse whole transcriptome analysis

Using nanoString’s GeoMx digital spatial profiling (DSP), we performed multiplexed and spatially resolved profiling analysis on FFPE samples. Briefly, 5 µm-thick FFPE tissue section was mounted on a glass slide for nanoString’s GeoMx digital spatial profiling (DSP) study. After deparaffinization, tissue was incubated in Tris-EDTA buffer (pH.9.0) in a steamer for 20 min for target retrieval, followed by incubation with PBS buffer containing 1 ug/mL of proteinase K at 37° C for 15 min for RNA target exposure. In Situ Hybridization was conducted by incubating the tissue section with RNA probe mixtures (GeoMs WTA Mouse RNA Probes for NGS) at 37° C overnight, and the tissue section was then stained at room temperature for one hour with fluorescence-conjugated morphology markers antibodies. We used anti-GFAP Alexa Fluor^TM^488 (0.5mg/mL, Invitrogen # 53-9892-82), anti-NeuN Alexa Fluor^TM^647 (1:100, EMD Millipore # ABN78), and nuclear dye of SYTO^TM^83 dye (250 nM, Thermofisher # S11364). Based on fluorescence imaging, regions of interest (ROIs) (200 µm–600 µm in diameter) were chosen for multiplexed and spatially resolved profiling analysis (3 ROIs/tumor). Photocleaved RNA probe oligos were transferred into a microwell and sequenced using next-generation sequencing (NGS) Illumina workflows. After the NGS readout, data were processed using an automated data processing pipeline to convert FastQ files to GeoMx-compatible DCC files. The DCC files were used for further analysis using nanoString’s inbuilt software.

Q3 normalized data was used to perform the downstream analysis. First, principal component analysis was performed on the segment-level data, and 4 outliers were identified (ROIs): 013 287325- T1, 002 289017-T2, 007 288997-T1, and 002 289037-T2. Then, the average expression measurement of segments per sample was calculated for each gene after filtering out the 4 outliers identified above (ROIs). Differential expression analysis was performed using the sample average values. Wilcoxon tests were performed between samples in X-ray exposed mice and controls, ^56^Fe ion exposed mice and controls, and ^56^Fe ion exposed mice and X-ray exposed mice. The “fdr” method in p.adjust function in R was utilized to adjust p-values for multiple comparisons. The cut-off for declaring significant differentially expressed genes between groups was p-value < 0.05. Finally, the mouse Hallmark gene sets (mh.all.v0.3) were obtained from the Molecular Signatures Database [19]. Geneset enrichment analysis was done using the R fgsea package with default parameters.

### Statistical analysis

We used sex-balanced and age-matched littermate controls in all experiments. After Adeno-Cre intranasal inhalation and whole-body irradiation in mice, the investigators were blinded to the type of radiation exposure at the time of complete necropsy and data collection. Adenoma/carcinoma-free, esthesioneuroblastoma (ENB)-free, and lymphoma-free survival was defined using histopathology analysis of tumor samples collected at the end of 1-year follow-up. Bar graphs and Kaplan-Meier plots were generated using GraphPad Prism (version 9.5.0), and significance was determined by log-rank tests and chi-square tests, respectively. In Volcano plots, each dot represented a gene with -log10(p-value), and significance was calculated using the Wilcoxon test. The heatmap shows log2 gene expression of all the significantly differentially expressed genes (DEGs) (Control vs. X-rays, Control vs. ^56^Fe ions, and X-rays vs. ^56^Fe ions comparisons). Venn diagrams showed the number of overlapping DEGs that were up-or down-regulated in Control vs. X-rays and Control vs. ^56^Fe ions. For the mouse hallmark pathway analysis, we plotted normalized enrichment scores for molecular pathways with padj < 0.05. The mouse whole transcriptome analysis was performed by QuantBio LLC.

## Results

### Exposure to X-rays and ^56^Fe ions IR accelerate tumor development in mice

To determine if radiation quality and sex differences affect radiation-induced lung tumorigenesis in mice, we performed single fraction whole-body irradiation in male and female *Rb^fl/fl^; Trp53^fl/+^* mice using different doses of 320 kVp X-rays (low-LET IR) or 600 MeV/n ^56^Fe ions (high-LET IR) and followed mice for tumor development for up to 2 years (Supplementary Figure 1A). Roughly two weeks before radiation exposure, the mice were treated with intranasal inhalation of Adeno-Cre to delete both copies of the *Rb* gene and a single copy of the *Trp53* gene. As expected, we observed a high prevalence of lung tumor formation in control, low-LET IR, and high-LET IR exposed mice (Supplementary figure 1H, 1I). However, we also observed a high prevalence of lymphomas and esthesioneuroblastomas (ENB) across groups. We observed that 4 Gy low-LET IR significantly accelerated lung tumors, lymphomas, and ENB compared to control mice without IR (Supplementary figures 1B-1D). Interestingly, we found a significantly higher incidence of lung and ENB tumors, but not lymphomas in high-LET IR exposed mice compared to control mice without IR (Supplementary figures 1E-1G). Overall, the lung tumors were the most common malignancies in X-ray and ^56^Fe ion IR cohorts (Supplementary figure 1H, 1I). The second most frequent malignancies were lymphomas and ENBs in X-rays and ^56^Fe ion cohorts, respectively (Supplementary figure 1H, 1I). Unfortunately, many mice were found dead overnight or on weekends, limiting the collection of tissues for histopathology confirmation. In addition, as mice were followed for different lengths of time, mice that died from one type of tumor may not have lived long enough to develop a different tumor. As a result, it was difficult to assess the impact of sex differences on low-and high-LET radiation-induced tumorigenesis in mice.

To address these limitations of the previous experiment, we designed a more controlled experiment in which we sacrificed mice approximately one year after radiation exposure (300-350 days), regardless of signs of malignancy, and performed a complete necropsy to collect all major organs for histopathology analysis (Figure 1A). Second, we kept one irradiation dose common between X-rays and ^56^Fe ions for direct comparison of low-and high-LET radiation-induced tumorigenesis. As shown in Figure 1B, lower doses of low-LET IR did not significantly impact tumor-free survival, but 4 Gy low-LET IR significantly reduced tumor-free survival compared to the 0 Gy cohort. We also noticed that multiple primary malignancies (MPM) or metastasis-free survival was significantly lower in the 4 Gy X-ray compared to the 0 Gy cohort (Figure 1C). Histopathology analysis revealed that lung adenoma/carcinoma and lymphomas were the most frequent malignancies in mice exposed to X-rays (Figure 1D). We detected that even lower doses of ^56^Fe ions IR (0.5 and 1 Gy) were sufficient to significantly reduce malignancy-free survival compared to the 0 Gy cohort (Figure 1E). In addition, 1 Gy ^56^Fe ions IR caused significantly reduced MPM or metastasis-free survival compared to the 0 Gy cohort (Figure 1F). In contrast to low-LET X-rays, esthesioneuroblastomas (ENB) were the predominantly detected malignancy in mice exposed to high-LET radiation (Figure 1G). Furthermore, we found rare malignancies, such as mammary carcinoma, harderian gland adenoma, granulosa cell tumor of the ovary, hepatoblastoma, pituitary adenoma, and small intestinal adenoma, post low-and high-LET radiation exposure (Supplementary figure 2). Overall, the data suggest that lower doses of high-LET IR significantly accelerate lung and ENB tumorigenesis in mice compared with low-LET IR.

**Figure 1:**
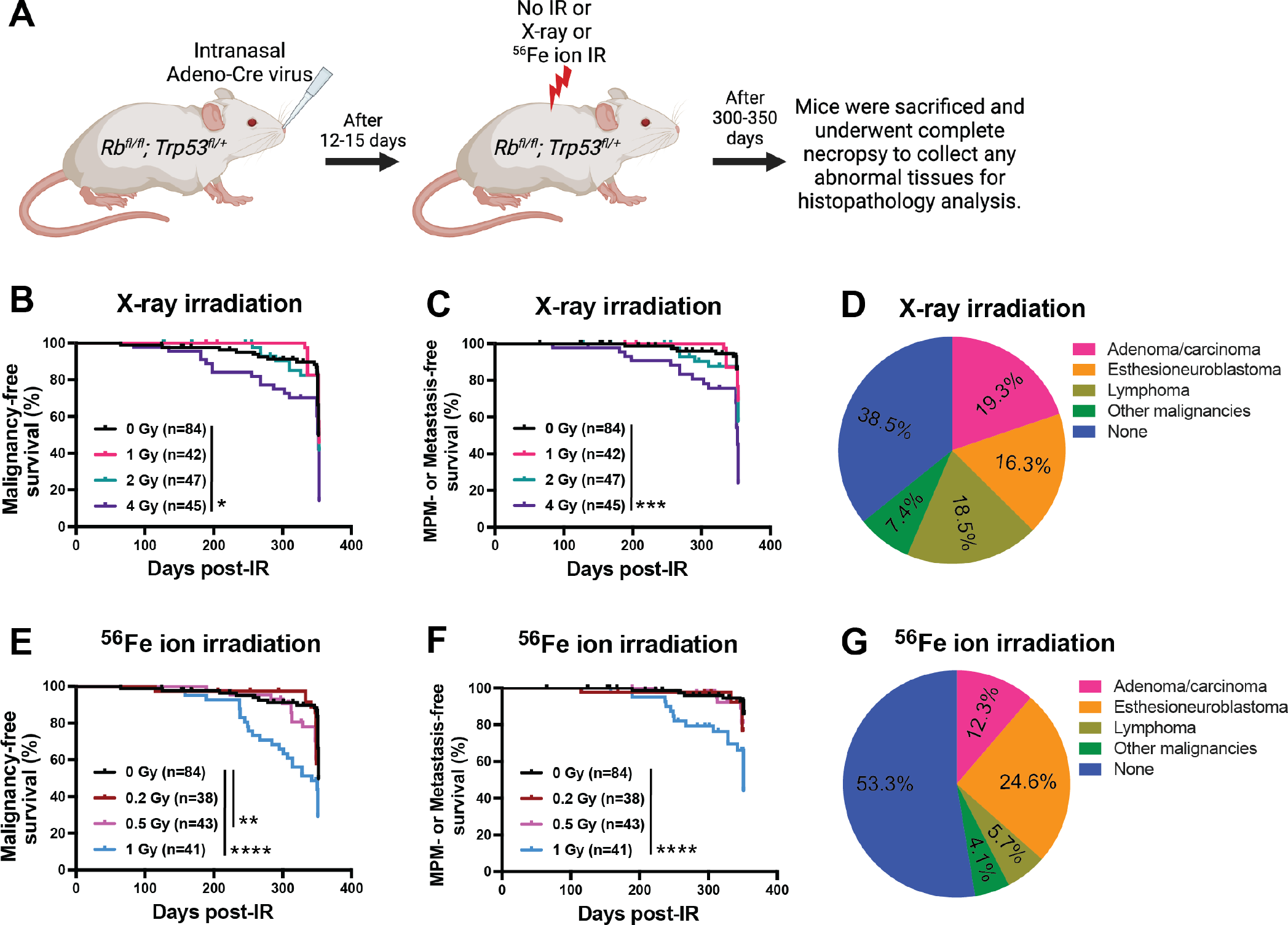
Incidence of tumorigenesis post whole-body exposure to X-rays versus ^56^Fe ions in *Rb^fl/fl^; Trp53^fl/+^* mice. (A) Schematic representation of experimental design highlighting timeline for intranasal injection of Adeno-Cre virus, radiation exposure, and follow-up. Kaplan-Meier plots show (B) Malignancy-free survival and (C) Multiple primary malignancies (MPM) or metastasis-free survival post whole-body 0, 1, 2, or 4 Gy X-ray exposure. (D) Pie graph shows the percentage incidence of adenoma/carcinoma, esthesioneuroblastoma, lymphoma, and other malignancies post whole-body X-ray exposure. Kaplan-Meier plots show (E) Malignancy-free survival and (F) Multiple primary malignancies (MPM) or metastasis-free survival post whole-body 0, 0.2, 0.5, or 1 Gy of ^56^Fe ion exposure. (G) Pie graph shows the percentage incidence of adenoma/carcinoma, esthesioneuroblastoma, lymphoma, and other malignancies post whole-body ^56^Fe ion exposure. n = number of mice. p-values in Kaplan-Meier plots were calculated using Log-rank (Mantel-Cox) tests. * = p<0.05, ** = p<0.01, *** = p<0.001, and **** = p<0.0001.

### Similar rate of overall malignancy development between male and female mice, regardless of the radiation quality

According to the Radiation Effects Research Foundation in Hiroshima, females exposed to radiation have a higher excess relative risk of solid cancer, including lung cancer [5]. Therefore, we assessed the impact of sex differences on the incidence of tumorigenesis post low-and high-LET radiation exposure. An increasing rate of overall malignancies was directly correlated with increasing doses of X-rays in male and female mice (Figure 2A). In addition, we found a similar rate of lung adenoma/carcinoma and ENB in male and female mice after whole-body X-rays (Figure 2B, 2C). However, it’s worth noting that female mice had a lower lung adenoma/carcinoma rate at baseline (0 Gy) compared to male mice, but this was not statistically significant. Similar to x-rays, we observed an increasing rate of overall malignancies with increasing doses of ^56^Fe ion exposure in male and female mice (Figure 2D). Interestingly, female mice had fewer overall malignancies compared to male mice after ^56^Fe ion exposure, but statistical significance was only observed in the 0.2 Gy cohort (Figure 2D). Similarly, female mice had a trend towards a lower rate of lung adenoma/carcinoma and ENB compared to male mice after ^56^Fe ion exposure, but the statistical analysis did not show any significant differences at any irradiation doses (Figures 2E, 2F). Thus, our data suggest that female mice are not at higher risk of solid tumorigenesis compared to male mice after exposure to whole-body X-rays or ^56^Fe ions.

**Figure 2:**
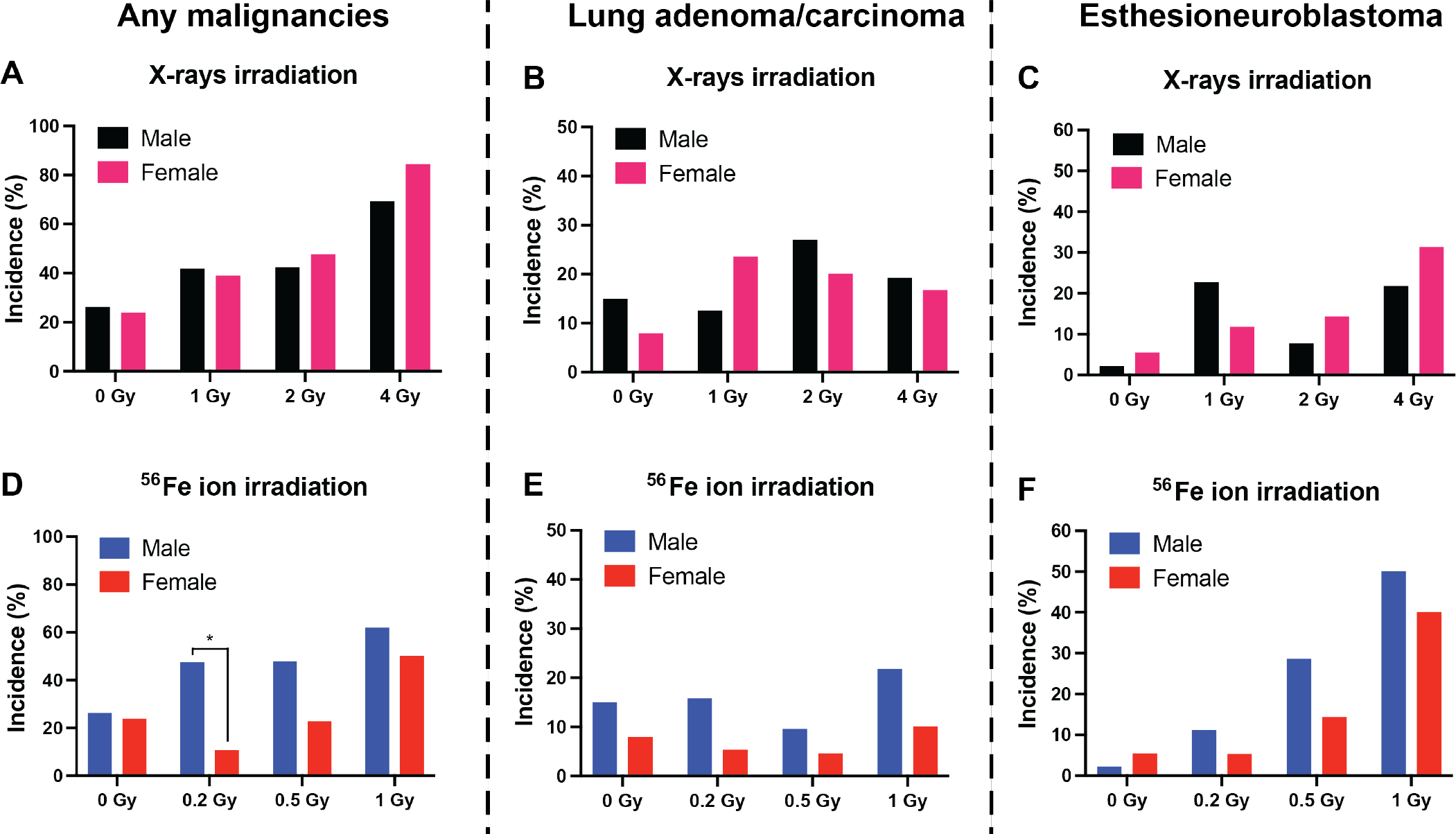
Incidence of tumorigenesis among male and female mice from second cohorts of mice. Bar graphs show the percentage incidence of (A) Any malignancies, (B) Lung adenoma/carcinoma, and (C) Esthesioneuroblastoma post whole-body 0, 1, 2, or 4 Gy of X-ray exposure. Bar graphs show the percentage incidence of (D) Any malignancies, (E) Lung adenoma/carcinoma, and (F) Esthesioneuroblastoma post whole-body 0, 0.2, 0.5, or 1 Gy of ^56^Fe ion exposure. p-values in all bar graphs were calculated using chi-square tests by comparing the incidence of malignancies between males and females at each radiation dose. * = p<0.05, ** = p<0.01, *** = p<0.001, and **** = p<0.0001.

### Significantly higher incidence of lung and brain malignancies post 1 Gy ^56^Fe ion IR compared to 1 Gy X-rays

As mentioned previously, lung adenoma/carcinomas, ENBs, and lymphomas were the most common malignancies whose development was accelerated by X-rays or ^56^Fe ions exposure. In addition, several preclinical studies have shown that high-LET radiation exposure has a higher relative biological effectiveness (RBE) compared to low-LET X-rays or *γ*-rays. For instance, ^56^Fe ion exposure causes a significantly higher incidence of lung cancers in C57BL/6 mice [20], liver cancers in C3H/HeNCrl mice [21], intestinal cancers in *APC^Min/+^* and *APC^1638N/+^* C57BL/6 mice [22], and lymphomagenesis in *Mlh1^+/-^* B6.129 mice [23] with RBE between 5-50 compared to low-LET X-ray or *γ*- ray exposure. The primary endpoint of our study was rate of tumorigenesis, and since a majority of malignancies were detected at the time of histopathology, we could not calculate RBE using a malignancy-free survival rate. Thus, we individually assessed and compared low-and high-LET-induced lung malignancies, ENBs, and lymphomas, followed by directly comparing the 1 Gy X-ray to 1 Gy ^56^Fe ion cohorts. First, we assessed radiation-induced lymphomas and discovered that the majority of lymphomas originated in the thymus and were most often disseminated to the liver or kidney (Supplementary figure 3A-3C). The rate of radiation-induced thymic lymphoma was low regardless of the type of radiation received. Only the 4 Gy X-ray cohort showed a significantly higher lymphomagenesis rate than the 0 Gy cohort (Supplementary figure 3D). None of the other irradiated cohorts exposed to X-rays or ^56^Fe ions showed a significantly higher lymphomagenesis rate than the 0 Gy cohort (Supplementary figure 3D-3F).

Next, we assessed the rate of lung malignancies and discovered that adenoma/carcinomas were the most common abnormalities found in the lungs post-radiation exposure (Figure 3A, 3B). None of the X-ray irradiated cohorts had a significantly higher rate of lung malignancies compared to the 0 Gy cohort (Figure 3C). 1 Gy ^56^Fe ions, but not 0.5 Gy or 0.2 Gy ^56^Fe ions, showed a significantly higher rate of lung adenoma/carcinoma compared to the 0 Gy cohort (Figure 3D). Furthermore, when we compared 1 Gy X-rays to 1 Gy ^56^Fe ions, we discovered that exposure to ^56^Fe ions was more efficient in accelerating lung malignancies over X-rays (Figure 3E). As shown in Figure 1, ENB was the primary malignancy in ^56^Fe ion exposed mice, so next, we assessed the rate at which X-rays and ^56^Fe ions induced brain malignancies in mice. Histopathology analysis showed that ENB was the most common brain malignancy found in irradiated cohorts and most often disseminated to the liver and lymph nodes (Figure 2F-2H). Similar to lung malignancies, none of the X-ray cohorts showed a significant increase in the rate of ENB compared to the 0 Gy cohort (Figure 3I). In contrast, ^56^Fe ion exposure caused a significantly higher incidence of ENBs post 0.5 Gy and 1 Gy irradiation compared to 0 Gy (Figure 3J). In addition, 1 Gy ^56^Fe ion exposure was more efficient in accelerating ENB over X-rays (Figure 3K). Our results demonstrate that the RBE for high-LET IR is significantly greater than 1.

**Figure 3:**
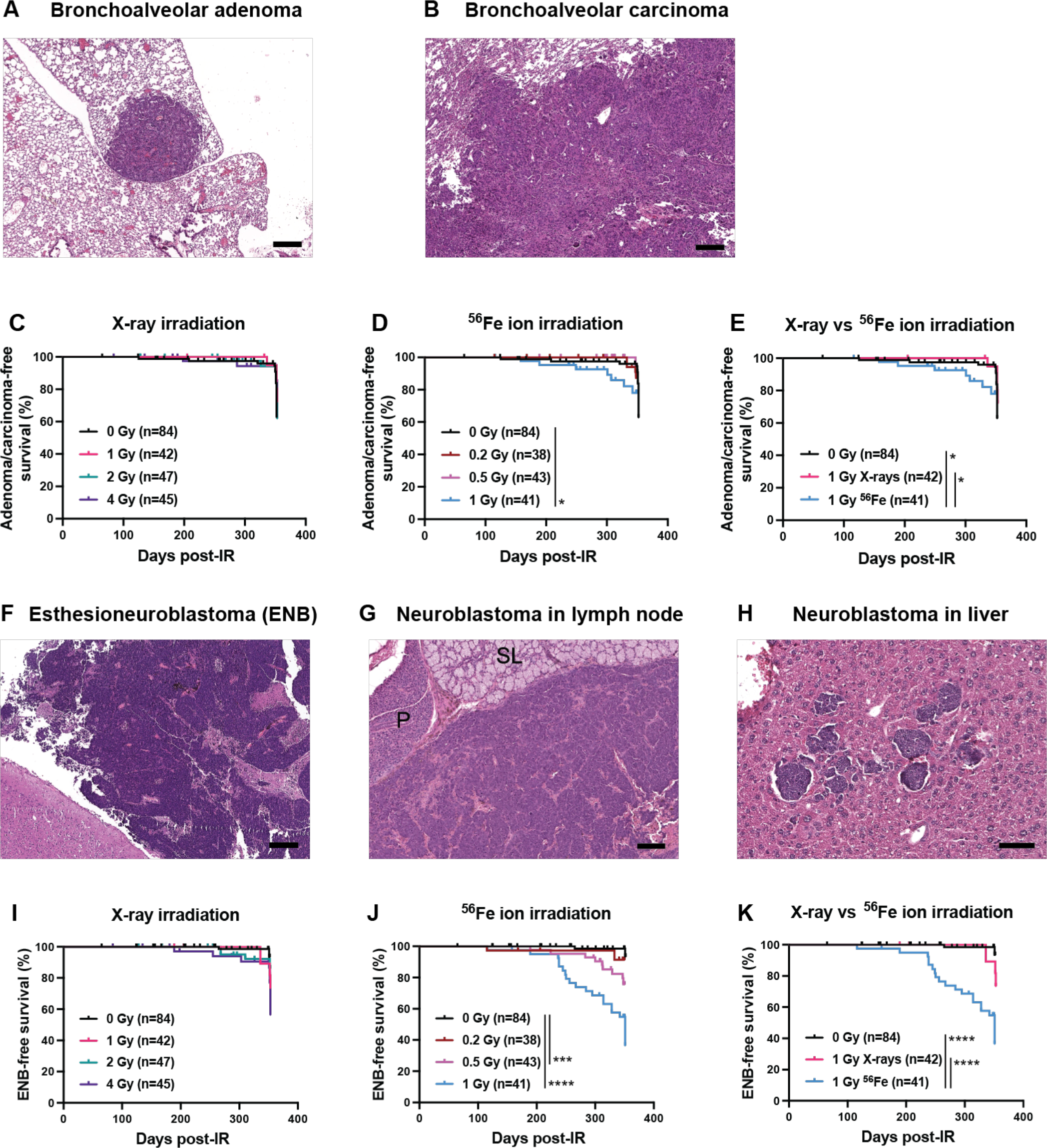
Incidence of bronchoalveolar adenoma/carcinoma and Esthesioneuroblastoma (ENB) in unirradiated and irradiated cohorts. (A) Discrete, well-circumscribed bronchoalveolar adenoma (4X, bar = 200um). (B) Poorly circumscribed and infiltrative bronchoalveolar carcinoma is characterized by irregular lobules and clusters of neoplastic cells invading adjacent pulmonary tissue (4X, bar = 200um). Kaplan-Meier plots show bronchoalveolar adenoma/carcinoma-free survival post whole-body (C) 0, 1, 2, or 4 Gy X-ray exposure, (D) 0, 0.2, 0.5, or 1 Gy ^56^Fe ion exposure, and (E) 1 Gy X-ray versus 1 Gy ^56^Fe ion exposure. (F) Large, invasive esthesioneuroblastoma effacing the olfactory bulb of the brain (4X, bar = 200um). (G) Olfactory neuroblastoma metastasis effacing the submandibular lymph node and compressing adjacent parotid (P) and sublingual (SL) salivary glands (4X, bar = 200um). (H) Metastatic clusters of olfactory neuroblastoma within liver sections (10X, bar =100um). Kaplan-Meier plots show the incidence of ENB-free survival post whole-body (I) 0, 1, 2, or 4 Gy X-ray exposure, (J) 0, 0.2, 0.5, or 1 Gy ^56^Fe ion exposure, and (K) 1 Gy X-ray versus 1 Gy ^56^Fe ion exposure. n = number of mice. p-values in Kaplan-Meier plots were calculated using Log-rank (Mantel-Cox) tests. * = p<0.05, ** = p<0.01, *** = p<0.001, and **** = p<0.0001.

### Distinct gene expression profiles but similar hallmark pathway enrichment between X-ray and **^56^**Fe ion-induced ENB

The incidence of ENB was significantly higher post ^56^Fe ions compared to X-rays. Therefore, we utilized whole transcriptome analysis of ENBs to identify potential differences in molecular pathway(s) responsible for control, X-ray-induced, and ^56^Fe ion-induced ENBs. Using nanoString’s GeoMx Mouse Whole Transcriptome Assay, we performed multiplexed & spatially resolved profiling analysis using 12 ENB formalin-fixed paraffin-embedded samples from three different cohorts (3 control, 5 X-ray and 4 ^56^Fe ion). First, we performed H&E staining followed by immunofluorescence staining of the same samples (serial sections) utilizing fluorescence-conjugated morphology marker antibodies. The fluorescence-conjugated morphology markers were used to identify regions of interest (ROIs) within the tumor for ROI selection. As shown in Figure 4A, ENB illuminated with green fluorescent color due to Syto^TM^83 binding to nucleic acids in tightly packed tumor cells. After ROI selection, downstream processing for multiplexed & spatially resolved profiling analysis showed over 1000 genes differentially expressed among the three cohorts of ENBs. Unsupervised clustering of differentially expressed genes revealed that IR-induced ENBs clustered separately from the control (unirradiated) cohort (Figure 4B). Within the IR-induced cohorts, ^56^Fe ion-induced ENBs clustered together, and most genes were down-regulated in ^56^Fe ion-induced compared to X-ray-induced ENBs (Figure 4B). Next, we individually compared X-ray and ^56^Fe ion cohorts to the control cohort to identify significantly altered genes that could explain the high rate of ENB in irradiated cohorts and potentially explain the differences between low-versus high-LET-induced ENBs. The gene expression profile of X-ray and ^56^Fe ion cohorts were distinct, with a higher number of genes differentially expressed in ^56^Fe ion-induced (336 genes) compared to X-ray-induced ENBs (200 genes) (Figure 4C-4D). As shown by Venn diagrams (Figure 4E), only 25 genes were found to be mutually differentially expressed among X-ray and ^56^Fe ion cohorts, suggesting that low-and high-LET irradiation exposure cause distinct gene expression signatures. To gain insight into significantly altered molecular pathways in each cohort, we performed Geneset Enrichment Analysis using differentially expressed genes to identify which hallmark pathways were preferentially altered between X-ray and ^56^Fe ion-induced ENBs. We discovered that MYC targets, oxidative phosphorylation, and MTORC1 signaling pathways were the most enriched in the irradiated cohorts, while the KRAS signaling pathway was the most enriched in the control cohort (Figure 4F-4G). The hallmark pathway analysis showed that the most significantly altered molecular pathways were similar between X-ray and ^56^Fe ion-induced ENBs, suggesting that the underlying mechanism(s) driving ENB might be independent of radiation quality.

**Figure 4:**
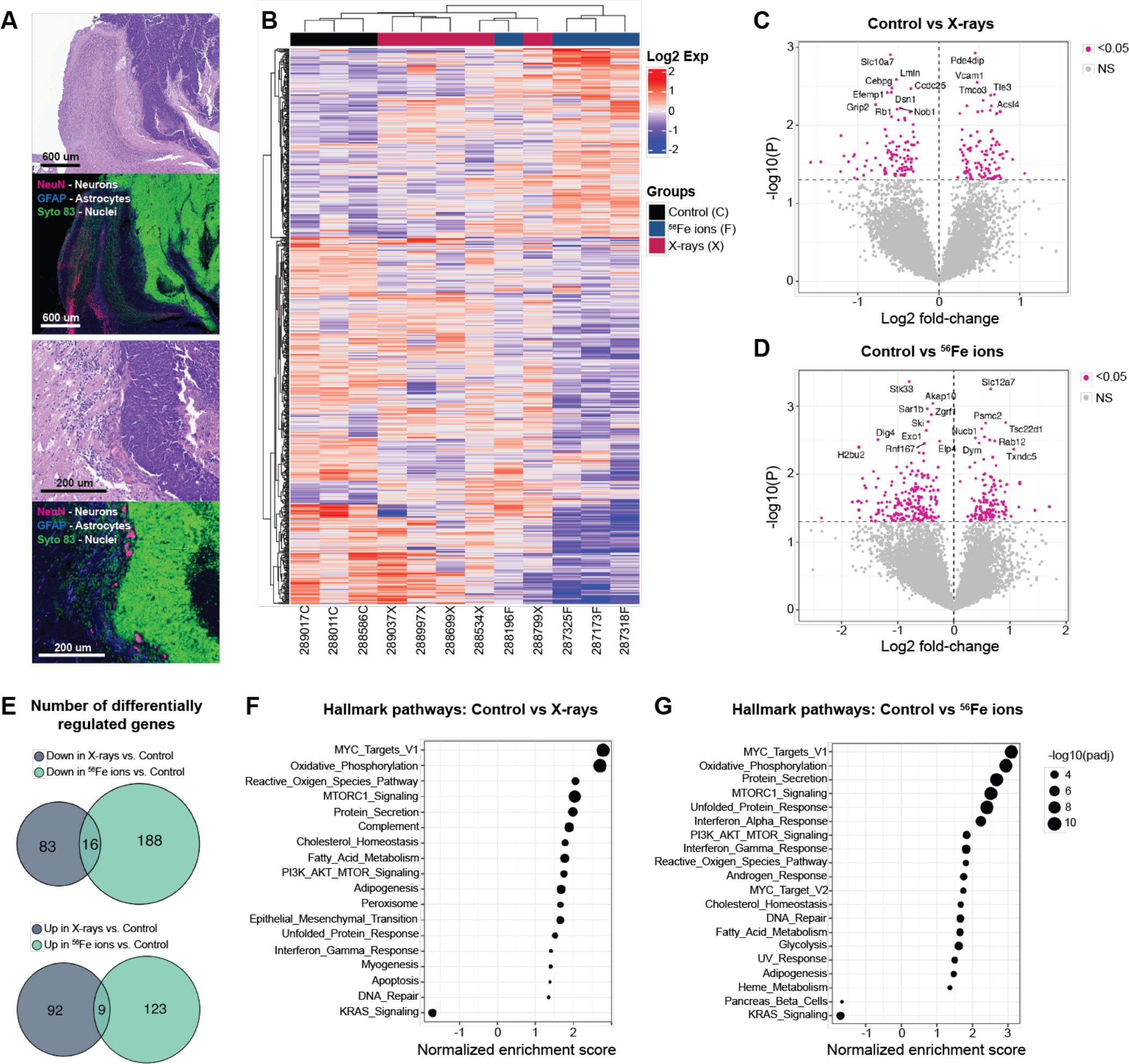
Esthesioneuroblastoma (ENB) whole transcriptome analysis using nanoString’s GeoMx digital spatial profiling. (A) Representative H&E and immunofluorescence images of the same samples (serial sections) utilizing fluorescent antibodies against NeuN (neuronal marker), GFAP (Astrocyte marker), and Syto 83 (Nuclei/DNA marker). (B) Heatmap shows unsupervised clustering of differentially expressed genes in control, X-ray, and ^56^Fe ion cohorts of ENBs. Volcano plots show differentially expressed genes in (C) Control vs. X-ray-induced ENBs or (D) Control vs. ^56^Fe ion-induced ENBs. (E) Venn diagrams show a number of down- & up-regulated genes in X-ray-and ^56^Fe ion-induced ENBs compared to the control. Hallmark pathway analysis shows significantly altered molecular pathways in (F) Control vs. X-ray-induced ENBs or (G) Control vs. ^56^Fe ion-induced ENBs. In volcano plots, p-values associated with differentially expressed genes were found using the Wilcoxon test.

## Discussion

Despite the overall decrease in lung cancer incidence in the Western world, studies suggest that women are at higher risk of developing lung cancer compared to men [24-26], and the incidence of lung cancer, in particular adenocarcinoma, is magnified among never-smoker younger women [26-28]. In addition, there is direct evidence that radiation exposure increases lung cancer incidences in humans, especially in women. One of the best data sets for human radiation exposure and cancer risk comes from atomic bomb survivors, where a study found that women had more than a two-fold increase in risk compared to men for radiation-induced lung cancer mortality [29]. Recent mathematical modeling of radiation-induced lung cancer from the atomic bomb survivor data suggested a model with clonal expansion of premalignant cells and sequential mutagenesis that fits the lung cancer data for male and female survivors [30]. In addition, epidemiology studies suggest that most lung cancers develop after sequential mutations in as few as three genes [11], including *TP53* and *RB* tumor suppressors [16, 31, 32]. Thus, to appropriately model the middle-aged astronaut population, we used a genetically engineered mouse model in which complete deletion of *Rb* and partial deletion of *Trp53* were achieved before mice were irradiated with low-or high-LET irradiation. A recently published study showed that male and female B6C3F1 mice develop lung cancer malignancies (adenoma and adenocarcinoma) at a similar rate post *γ*-ray or carbon ion exposure [33]. Similarly, we observed that female mice were not at higher risk of radiation-induced lung malignancies compared to male mice, regardless of the radiation type. Generally, mice are more sensitive to radiation-induced cancers than humans, and the current study was performed using a single fraction of IR at a higher dose-rate. However, astronauts in outer space will be exposed to many types of HZE particles with a wide-range of energies and dose-rates. Therefore, future studies with a mixed ion beam field and lower dose-rate will be helpful to investigate the finding that female mice are not at higher risk of radiation-induced lung cancers than male mice.

It is an accepted fact by NASA that space radiation exposure will be a significant concern and limiting factor for long-term space missions in the future. Over the past few decades, *in vivo* studies have confirmed that RBE is significantly higher for high-LET radiation-induced carcinogenesis than low-LET irradiation for most malignancies [3]. Similarly, we confirmed that 1 Gy ^56^Fe ion exposure caused a significantly higher incidence of lung adenoma/carcinoma and ENB compared to 1 Gy of X-rays. In addition, we found that time to ENB onset was reduced post ^56^Fe ion exposure compared to X-rays. After bone marrow, the most common sites of neuroblastoma metastasis in humans are lymph nodes and the liver [34, 35]. Similarly, we found a significantly higher number of mice with evidence of neuroblastoma in lymph nodes and liver post low-and high-LET radiation exposure, suggesting possible metastasis from primary ENB. High-LET IR has been previously shown to cause acute brain toxicity leading to behavioral defects [36-38], and our data suggest that radiation-induced brain malignancies might also be a significant concern for NASA.

High-LET radiation exposure induces tumors in most tissue types and is a potent carcinogen compared to low-LET X-rays or *γ*-rays, mainly due to the clustered DNA damage induced by high-LET IR [3]. A study of radiation-induced lymphomagenesis showed that even though low-and high-LET radiation caused distinct mutational and transcriptional changes, the underlying mechanism of lymphomagenesis was similar between low-and high-LET IR [23, 39]. Another study has shown that mutational and structural changes at the genomic level are distinct in low-versus high-LET radiation-induced cancers in *Trp53* deficient mice [40]. We performed a whole transcriptome analysis to evaluate gene expression changes associated with X-ray versus ^56^Fe ion-induced ENB. Similar to previously published studies, we found that there was little similarity in gene expression patterns and commonly altered genes between X-ray and ^56^Fe ion-induced ENBs. In contrast, hallmark pathway analysis using differentially expressed genes revealed similar hallmark pathways altered in radiation-induced ENBs, regardless of the radiation quality factor. The Myc pathway was the most significantly enriched hallmark pathway in X-ray and ^56^Fe ion-induced ENBs, which is interesting because most human neuroblastomas have been shown to rely on MYC pathway alteration [41, 42]. Finally, tumor development in our mouse model depended on the complete deletion of *Rb* gene and partial deletion of *Trp53* gene. Studies have shown that cancers driven by loss of *Rb* are associated with increased activity of the PI3K/AKT/MTOR pathway [43-45]. Similarly, we observed MTORC1 and PI3K/AKT/MTOR signaling pathway upregulation in radiation-induced ENBs, indirectly confirming the loss of *Rb-driven* ENBs in mice.

In summary, our findings suggest that ^56^Fe ions more potently initiate lung tumors and esthesioneuroblastomas compared to X-rays, but the underlying mechanism(s) of tumorigenesis might be similar between low-and high-LET induced tumors. In contrast to data from atomic bomb survivors, we did not observe a higher rate of radiation-induced cancers in female mice.

## Acknowledgements

This work was funded by grants from NASA (NNX11AC60G) and the NCI (2R35 CA197616) to DGK. We thank Adam Rusek and his team at NASA Space Radiation Laboratory and Peter Guida at Brookhaven National Laboratory for their assistance with these experiments. We also thank the Duke University BioRepository & Precision Pathology Center (Duke BRPC – supported by P30CA014236) and the National Cancer Institute’s Cooperative Human Tissue Network (CHTN – supported at Duke University by UM1CA239755) for helping with the nanoString’s GeoMx digital spatial profiling experiment.

