## Supplemental Figures for "^56^Fe ion exposure increases the incidence of lung and brain tumors at a similar rate in male and female mice"

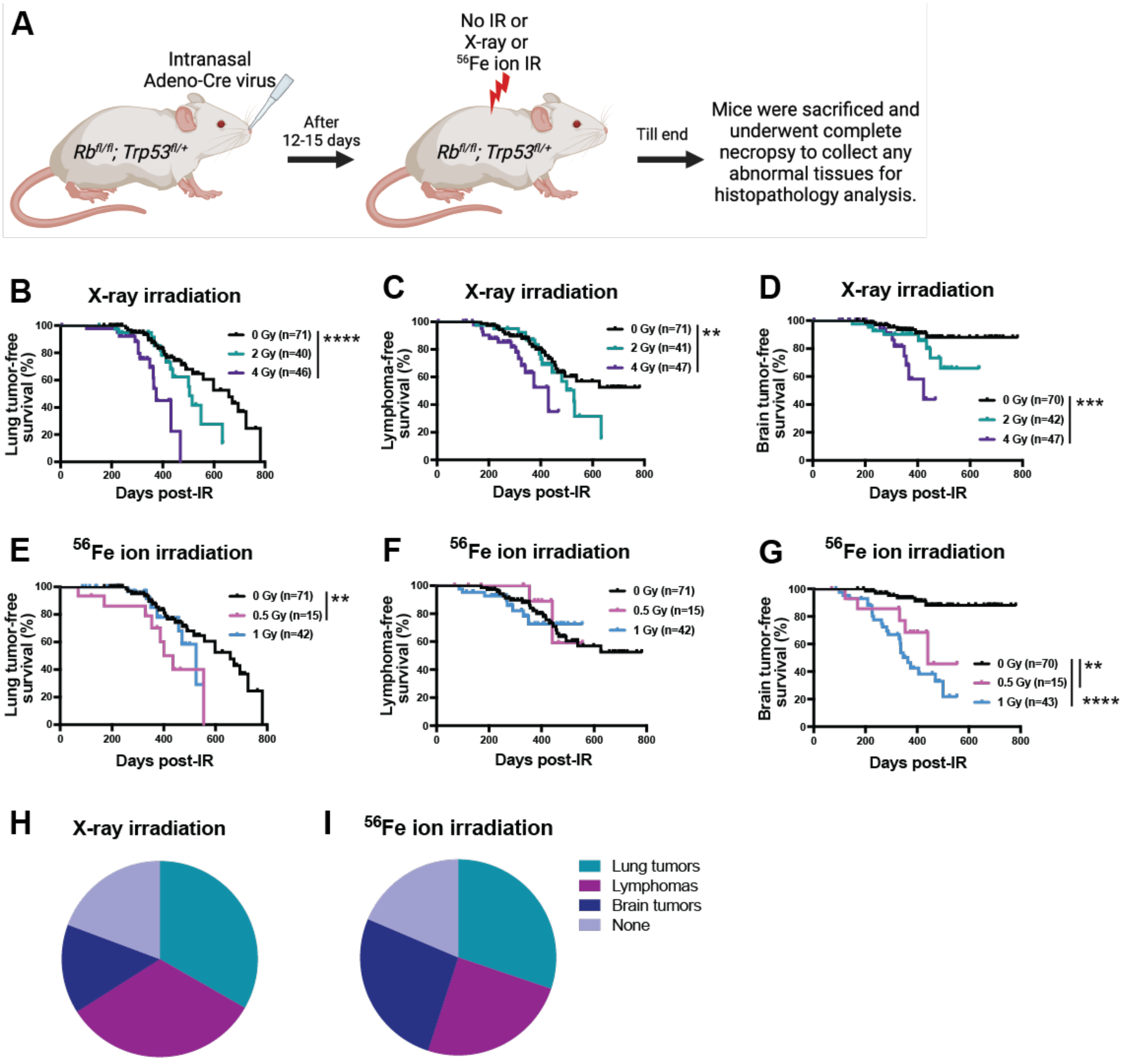


**Supplementary figure 1:** Incidence of tumorigenesis post whole body exposure to X-rays versus $^{56}Fe$ ions in *Rb*$^{fl/fl}$; *Trp53*$^{fl/+}$ mice. (A) Schematic representation of experimental design showing timeline for intranasal injection of Adeno-Cre virus, radiation exposure, and follow up. Kaplan Meier graphs show (B) Lung tumor-free survival, (C) Lymphoma-free survival, and (D) Brain tumor-free survival post whole body X-rays exposure. Kaplan Meier graphs show (E) Lung tumor-free survival, (F) Lymphoma-free survival, and (G) Brain tumor-free survival post whole body $^{56}Fe$ ions exposure. Pie graphs show the percentage incidence of lung tumors, lymphomas, and brain tumors in mice post whole body (H) X-ray (all doses combined) and (I) $^{56}Fe$ ion IR (all doses combined). n = number of mice. p-values in Kaplan Meier graphs were calculated by Log-rank (Mantel-Cox) tests. * = $p<0.05$, ** = $p<0.01$, *** = $p<0.001$, and **** = $p<0.0001$.

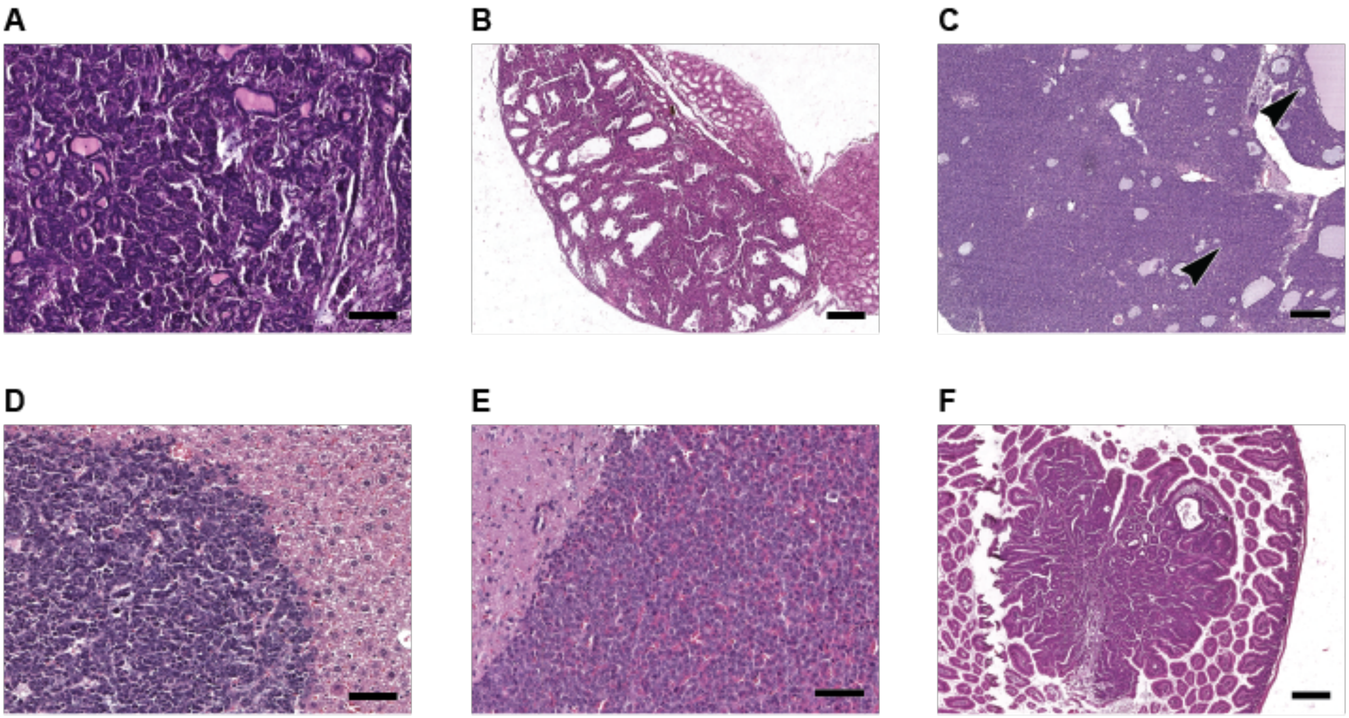


**Supplementary figure 2:** Other select neoplasms observed in *Rb*$^{fl/fl}$; *Trp53*$^{fl/+}$ mice. (A) Mammary carcinoma, composed of infiltrative tubules and clusters of neoplastic epithelial cells (10X, bar = 100um). (B) Harderian gland adenoma composed of lobules and tubules of well-differentiated epithelial cells, compressing adjacent normal Harderian gland (4X, bar = 200um). (C) Granulosa cell tumor of the ovary, characterized by sheets and lobules of neoplastic epithelium effacing normal ovarian structures, with remnants of ovarian follicles present (arrowheads, 4X, bar = 200um). (D) Hepatoblastoma in sections of liver, composed of dense basophilic neoplastic cells with angular nuclei and scant cytoplasm (20X, bar = 50um). (E) Pituitary adenoma composed of sheets of neoplastic cells compressing adjacent brain (20X, bar = 100um). (F) Small intestinal adenoma (mucosal polyp) along the mucosal surface of the duodenum (4X, bar = 200um).

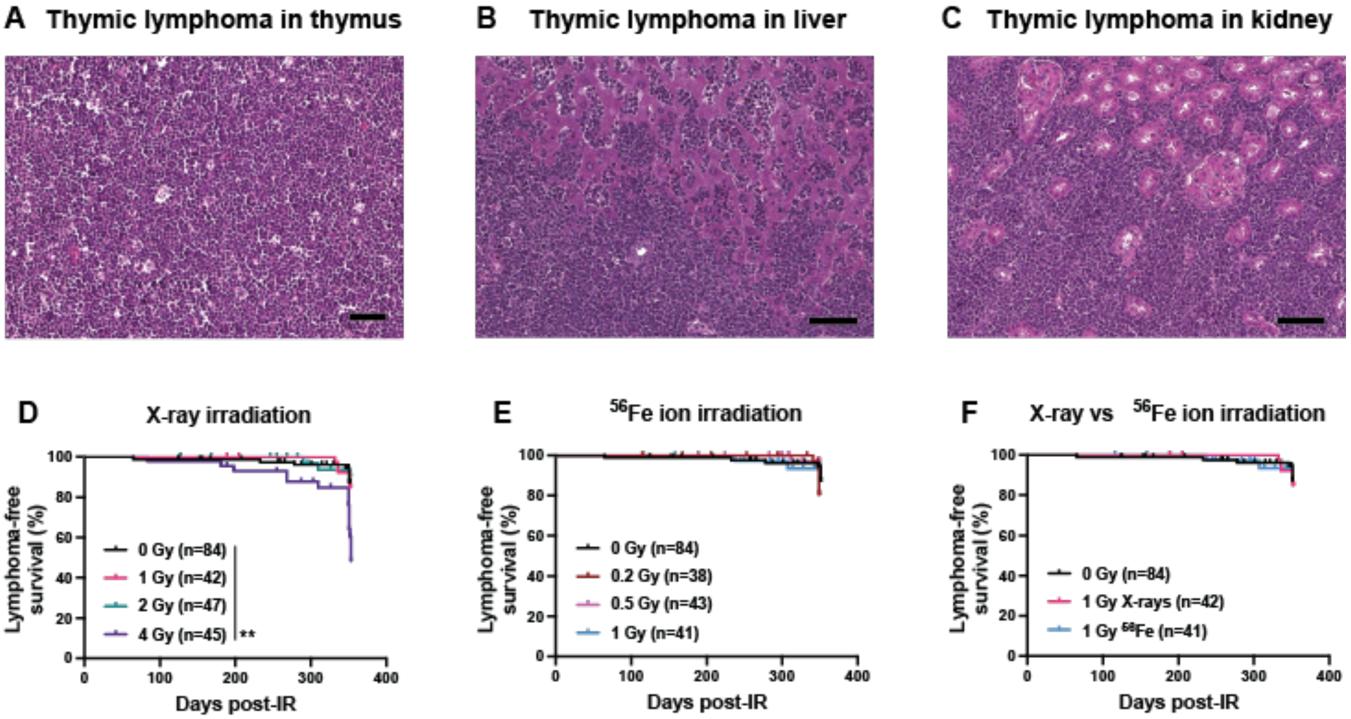


**Supplementary figure 3:** Incidence of lymphomagenesis post whole body exposure of X-rays versus $^{56}$Fe ions in *Rb*$^{fl/fl}$; *Trp53*$^{fl/+}$ mice. (A) Primary thymic lymphoma composed of sheets of neoplastic lymphocytes effacing the thymus (4X, bar = 200um). (B-C) Thymic lymphoma dissemination to the liver (B, 20X, bar = 50um) and kidney (C, 20X, bar = 50um). Kaplan Meier graphs show lymphoma-free survival post whole body exposure of (D) X-rays, (E) $^{56}$Fe ions, and (F) X-rays versus $^{56}$Fe ions. n = number of mice. p-values in Kaplan Meier graphs were calculated by Log-rank (Mantel-Cox) tests. * = p<0.05, ** = p<0.01, *** = p<0.001, and **** = p<0.0001.
